# Competition between continuously evolving lineages in asexual populations

**DOI:** 10.1101/062976

**Authors:** Noah Ribeck, Joseph S. Mulka, Luis Zaman, Brian D. Connelly, Richard E. Lenski

## Abstract

In an asexual population, the fate of a beneficial mutation depends on how its lineage competes against other mutant lineages in the population. With high beneficial mutation rates or large population sizes, competition between contending mutations is strong, and successful lineages can accumulate multiple mutations before any single one achieves fixation. Most current theory about asexual population dynamics either neglects this multiple-mutations regime or introduces simplifying assumptions that may not apply. Here, we develop a theoretical framework that describes the dynamics of adaptation and substitution over all mutation-rate regimes by conceptualizing the population as a collection of continuously adapting lineages. This model of “lineage interference” shows that each new mutant’s advantage over the rest of the population must be above a critical threshold in order to likely achieve fixation, and we derive a simple expression for that threshold. We apply this framework to examine the role of beneficial mutations with different effect sizes across the transition to the multiple-mutations regime.

## INTRODUCTION

Mutation and natural selection are two core processes in evolutionary theory, yet we lack a comprehensive theoretical framework for describing the dynamics of adaptation by natural selection as beneficial mutations sweep through asexual populations. Much of the difficulty arises from the *interference* between lineages with alternative contending beneficial mutations that compete with one another as they move toward (but often fail to achieve) fixation. Gerrish and Lenski developed a model of *clonal interference* based on the probability that a particular beneficial mutation will become fixed before a superior mutation outcompetes it (GERRISH AND LENSKI 1998). Their work has since gained wide acceptance in the field as the standard quantitative description of adaptation. However, clonal-interference theory rests upon a key assumption that each contending mutation arises on an identical genetic background. This assumption fails for populations that are sufficiently large or have high beneficial mutation rates, such that lineages accumulate multiple beneficial mutations before any single mutation achieves fixation (Fig. 1). Several papers have attempted to relax this assumption, but in so doing they introduced additional assumptions, such as constant fitness effects (ROUZINE *et al.* 2003; DESAI AND FISHER 2007; BRUNET *et al.* 2008; ROUZINE *et al.* 2008; HALLATSCHEK 2011). Thus, we still need an intuitive and quantitatively accurate framework that describes asexual adaptation in both the strong and weak mutation regimes.

**Fig. 1.**
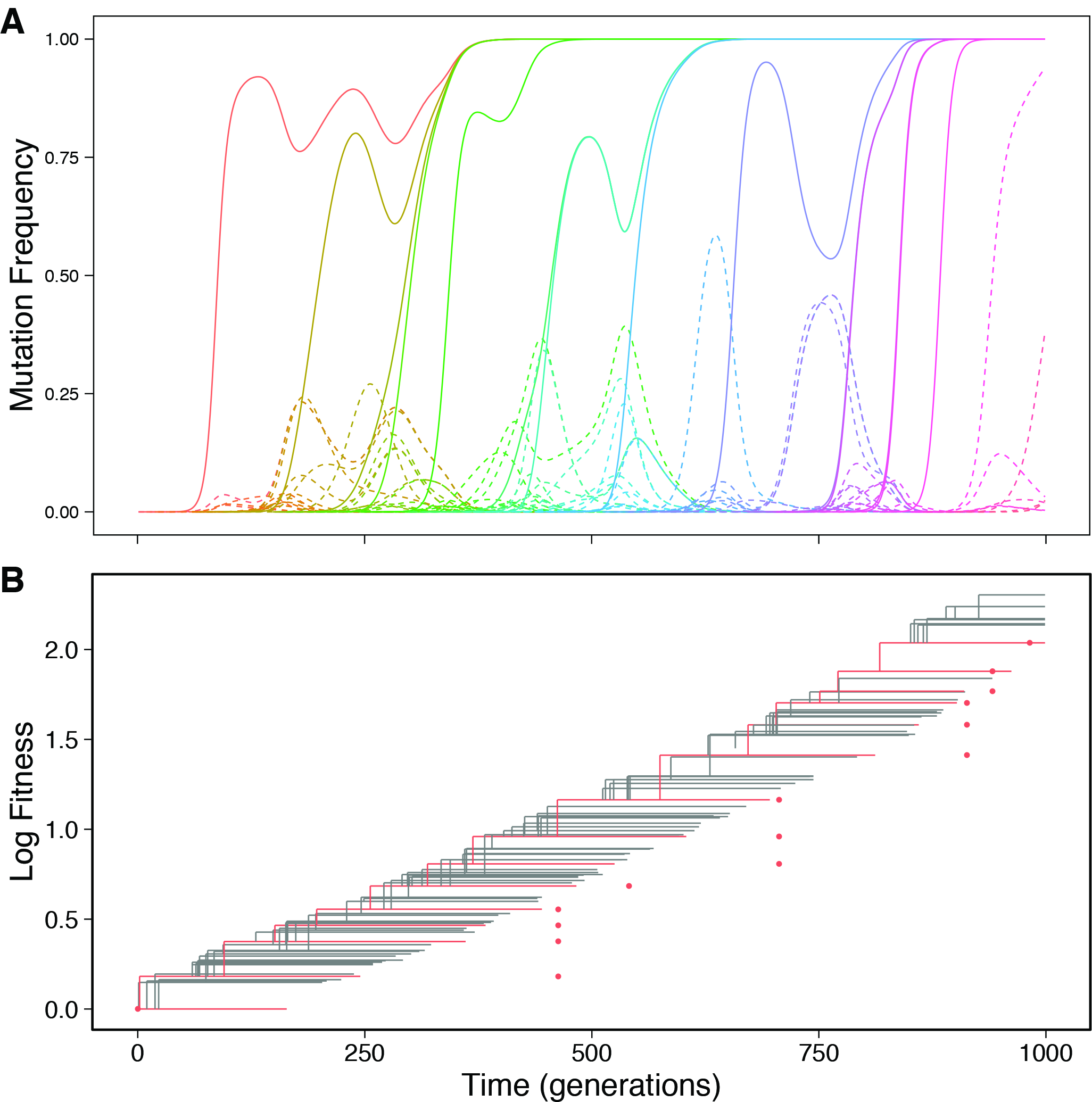
Concurrent sweeps and interference in the multiple-mutations regime. (A) Frequency dynamics of beneficial mutations in a simulated asexual population (see Methods) with size *N* = 10^7^, beneficial mutation rate *μ* = 10^−4^, and mean beneficial effect 1/*α* = 0.01. Mutations that eventually fix (solid lines) often occur on backgrounds with other beneficial mutations that have not yet fixed. There is also intense interference among contending lineages, often resulting in the extinction of beneficial mutations (dashed lines). (B) Corresponding phylogenetic tree. Red branches are genotypes where a beneficial mutation eventually fixes, at a time denoted by a red dot and a fitness increment indicated on the vertical axis. Often, a genotype is no longer extant when its original beneficial mutation becomes fixed, because it has since acquired additional beneficial mutations and its progenitor has been outcompeted.

Here, we develop a general population-genetic model for describing the dynamics of adaptation by natural selection in asexual populations, one that accounts for mutations arising on genetic backgrounds that already carry beneficial mutations that have not yet fixed. We will follow the mathematical logic of the clonal-interference model, but instead of describing the fates of *clones* in a population, we will describe the fates of *lineages*. In this model of *lineage interference*, we make the key observation that the expected rate of fitness increase for any lineage is proportional to its frequency in the population. Any segregating lineage must then have a large enough initial advantage over the rest of the population such that it reaches 50% frequency before its relative advantage is negated by adaptation of competing lineages. By comparison to simulations, we will show that this model provides a general description of asexual adaptation that accurately predicts the rates of adaptation and substitution across parameter regimes.

## METHODS

### Lineage-interference model

In clonal-interference theory, the likelihood of a particular beneficial mutation surviving interference is assessed by its probability of sweeping to fixation before any single superior mutation arises on the same background (GERRISH AND LENSKI 1998). In this formulation, each individual mutation competes for fixation against other individual mutations. This simplification breaks down, however, for populations with high *Nμ*, where *N*is the population size and *μ* is the beneficial mutation rate (per genome per generation).

To describe more accurately populations with a high mutation supply rate (high *Nμ*) — the *multiple-mutations regime* (DESAI AND FISHER 2007)—we introduce lineage-interference theory. This theory explicitly considers the genetic background in both the treatment of newly occurring mutations and the effect of interference on the fates of those mutations. First, the success of a new mutation depends on the sum of its beneficial fitness effect and its background fitness; for simplicity, we assume there is no epistasis. This “mutant fitness” determines an “effective” selective advantage that includes the effects of any pre-existing mutations carried by the genetic background. Second, the probability of fixation for a new mutation can be increased by its lineage’s acquisition of additional beneficial mutations, and decreased by beneficial mutations in competing lineages. In other words, the “mutant lineage” defined by the new mutation, having just acquired a bump in fitness, has a head start in a race against the rest of the population, which we will call the “pack lineage.”

As this race progresses, both lineages can acquire additional mutations, and the mutant lineage eventually either wins (achieves fixation) or loses (goes extinct). The original mutant will need a head start in order to win, because the pack lineage initially has many more opportunities to acquire additional beneficial mutations owing to its larger population size. To quantify how much of a head start the mutant lineage needs, we must express the adaptation rates for both the mutant lineage *v*_mut_ and the pack lineage *v*_pack_ that result from the accumulation of additional beneficial mutations and their spread by selection. In the absence of epistasis, both lineages have access to the same pool of additional beneficial mutations, and any additional mutation in either lineage is subject to the same potential effects of interference from other members of the population. Early mutations in the mutant lineage have an advantage by virtue of occurring on a higher fitness background: if the effect size of the second mutation is comparable to the original mutation, the double-mutant now has roughly twice the probability of surviving drift as the original. However, because the mutant lineage is initially outnumbered by a factor of *N*, the pack lineage has an *N*-times greater supply rate of new mutations, so *v*_mut_ is still negligible compared to *v*_pack_ when the mutant lineage is relatively new. Therefore, each lineage’s adaptation rate is approximately proportional to its frequency:

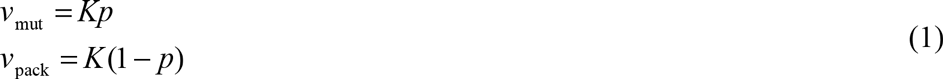

where *p* is the frequency of the mutant lineage and *K* is a proportionality constant. The adaptation rate *v* of the entire population, which is approximately constant in the absence of epistasis, is given by the weighted sum of the two groups:

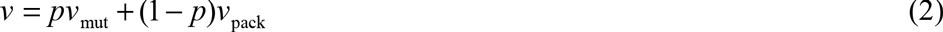

The mutant lineage has a relative fitness advantage *s* over the pack lineage—i.e., over the rest of the population. This advantage is measured relative to the mean fitness of the pack lineage, so it includes effects from the mutation itself and from its genetic background. The rate at which *s* changes is equal to the difference between the two groups’ respective adaptation rates:

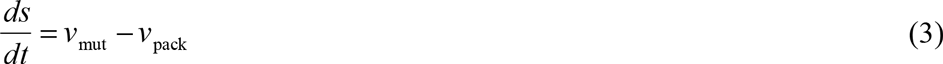

Together with Eqs. (1) and (2), this rate gives an expression for the changing relative advantage of the mutant lineage, in terms of constant *v* and time-dependent *P*:

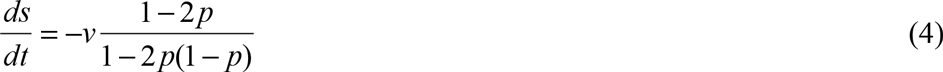

The changing relative frequency of the mutant lineage under continuous growth is given by (FISHER 1930) as:

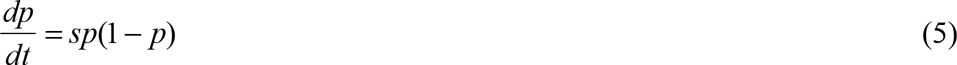

Eqs. (4) and (5) are the “lineage interference equations,” which provide a framework for determining the fate of the original mutant lineage. Because each lineage’s adaptation rate is proportional to its frequency, the mutant’s frequency must—if it is likely to survive—reach 50% before its mean fitness is surpassed by that of the pack lineage. This implies the existence of a critical threshold *s*_crit_ for the original mutant’s initial advantage *s*_0_, above which the original mutation is likely to fix, and below which it is likely to go extinct. Numerical solutions for these equations with different values of *s*_0_ show this critical behavior (see Fig. 2). The following section provides a derivation for an expression of *s*_crit_.

**Figure 2.**
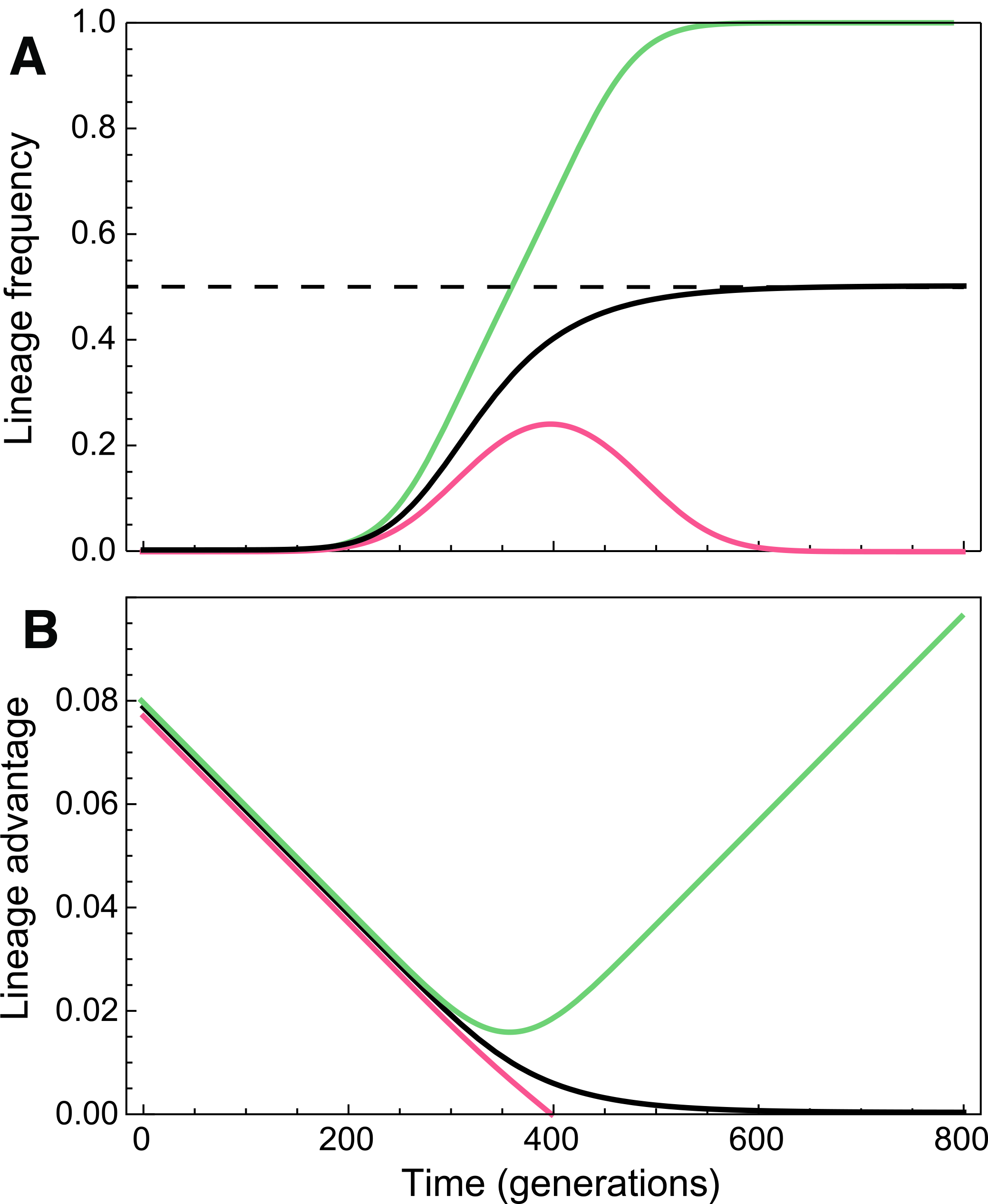
Lineage interference equations reveal critical threshold required for lineage fixation. Numerical solutions of the lineage-interference model (Eqs. 4-5) for time-dependent values of (A) frequency *p*, and (B) lineage advantage *s*; with initial advantage *s*_0_ = 0.077 < *s*_crit_ (red curves), *s*_0_ = 0.08 > *s*_crit_ (green curves), and *s*_0_ = 0.0784 = *s*_crit_ (black curves) where *s* is exactly depleted when *p* = 0.5. Population size *N* = 10^7^.

This framework is accurate when the mutant lineage is new and therefore rare, such that *v*_pack_ ≫ *v*_mut_ owing to the vastly greater mutation supply rate in the pack lineage. This assumption breaks down for later times when the two lineages are of more comparable frequency. However, we are specifically interested in cases near the critical threshold, where the mutant lineage’s advantage approaches zero when its frequency approaches 50%. Because the two lineages have comparable fitness at this point, the difference in their adaptation rates is still dominated by the mutation supply rate, so proportionality between adaptation rate and frequency still holds. Therefore, this framework is appropriate for analyzing the potential outcomes of a mutant lineage near the critical threshold.

### Minimum advantage required for fixation

The differential equation system defined by Eqs. (4) and (5) has no analytical solution, so we must seek an approximation for the critical threshold *s*_crit_ for a mutation to become fixed. First, consider the dynamics of a beneficial mutation without interference, assuming that it has escaped drift loss while rare. Such a mutation sweeps through a population of size *N* according to Eq. (5) with constant *s*, which has the solution:

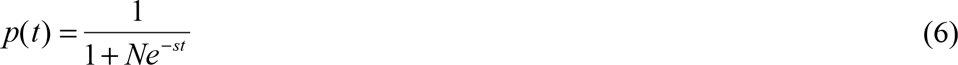

It takes log(*N*)/*s* generations for this mutation to reach 50% frequency. In the lineage-interference model, a mutant lineage near the critical threshold would take roughly twice this long to approach 50%, because its relative advantage spends most of its time decreasing nearly linearly toward *s* = 0 (Fig. 2) and therefore has a time-average advantage of *s*_crit_/2.

However, this approximation does not account for any further adaptation achieved by the mutant lineage itself, which occurs mostly at later times when it has reached a substantial frequency. To estimate this effect, consider the integral of *p* for a constant-s mutation up to the time required to achieve 50% frequency *t*_half_ = log(*N*)/*s*:

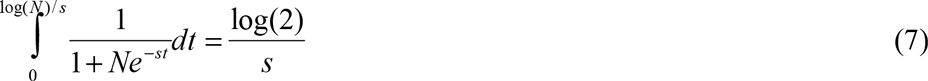

Each lineage’s adaptation rate is proportional to *p*, and so this integral is the cumulative increase in *s* that results from its ongoing adaptation. And because *t*_half_ is inversely proportional to *s*, this integral also represents the decrease in *t*_half_ caused by its adaptation. Together with the factor of 2 explained above, this process of ongoing adaptation gives an estimate of *t*_half_ for a mutant lineage near the critical threshold:

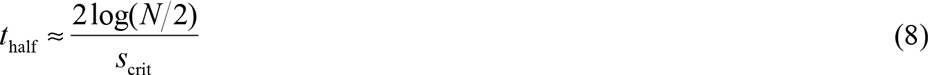

At this point, when the mutant lineage approaches 50% frequency, its initial advantage is reduced to zero, which takes roughly *s*_crit_/*v* generations. Solving for *s*_crit_, this gives:

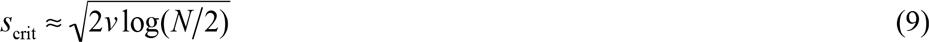

This critical value represents the minimum required fitness difference between a new mutant and the mean fitness of the rest of the population in order for the new mutation to eventually become fixed in a population of size *N* with an overall rate of adaptation *v*. Eq. (9) provides an excellent approximation for numerical solutions of Eqs. (4) and (5), as shown in Fig. 3; it does not depend on assumptions about the magnitude or distribution of mutational effects, although it breaks down for very small population sizes.

**Figure 3.**
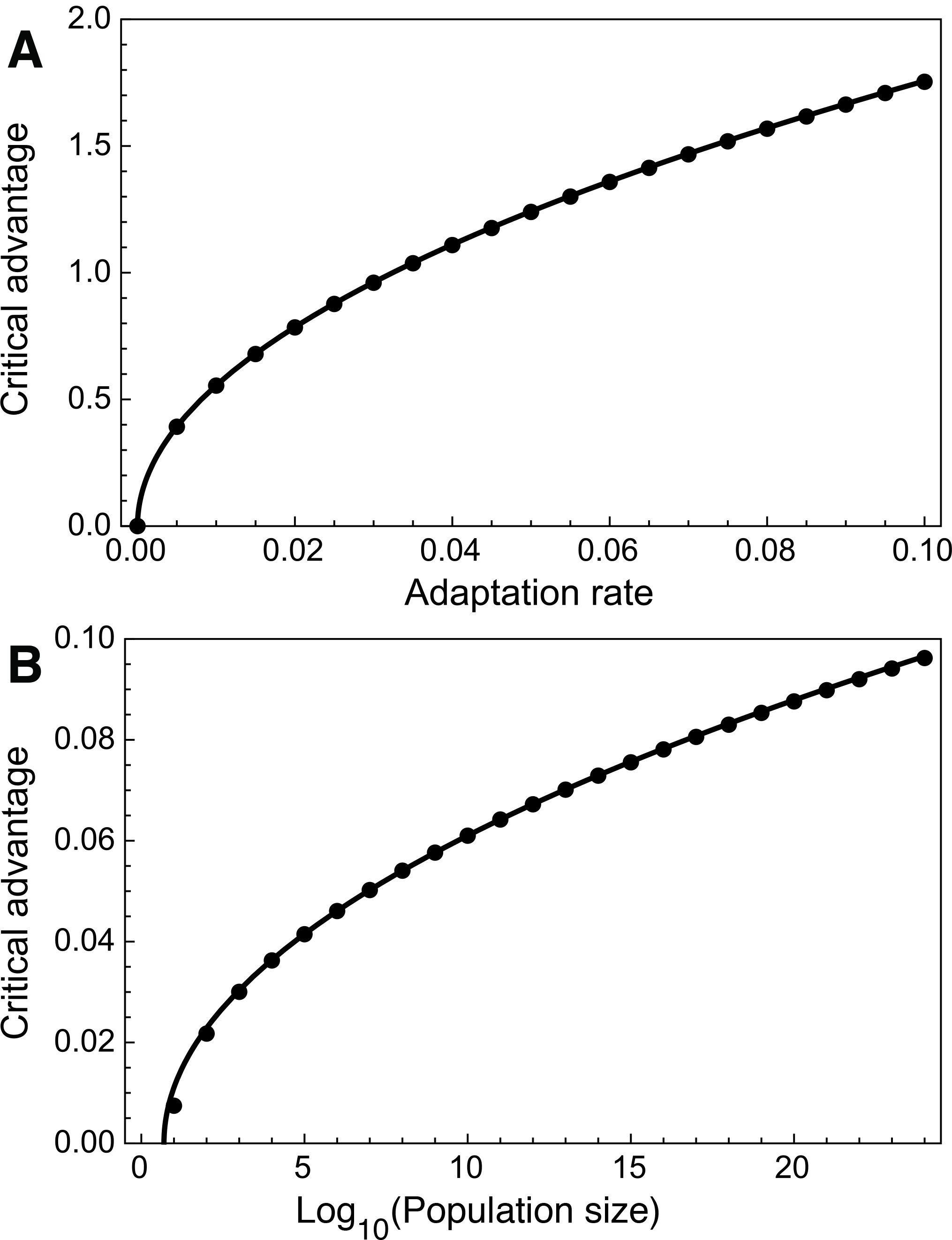
Closed-form approximation of the critical threshold for lineage fixation matches numerical solutions. Numerical solutions of *s*_crit_ from the lineage interference equations (points) closely match the closed-form approximation in Eq. (9) (curves) for both (A) varying adaptation rate *v* with *N* = 10^7^ (relative error ~0.17% on all points), and (B) varying population size *N* with *v* = 2 × 10 “^4^ (relative error ~0.3% on all points with *N*≥10^4^).

### Numerical methods and simulations

Numerical solutions throughout were found using Mathematica 10. The `NDSolve` function was used for the lineage interference equations (Eqs. (3) and (4), plotted in Fig. 3), and the `FindMinimum` function was used for the adaptation rate (Eq. (14), plotted in Fig. 4A).

**Figure 4.**
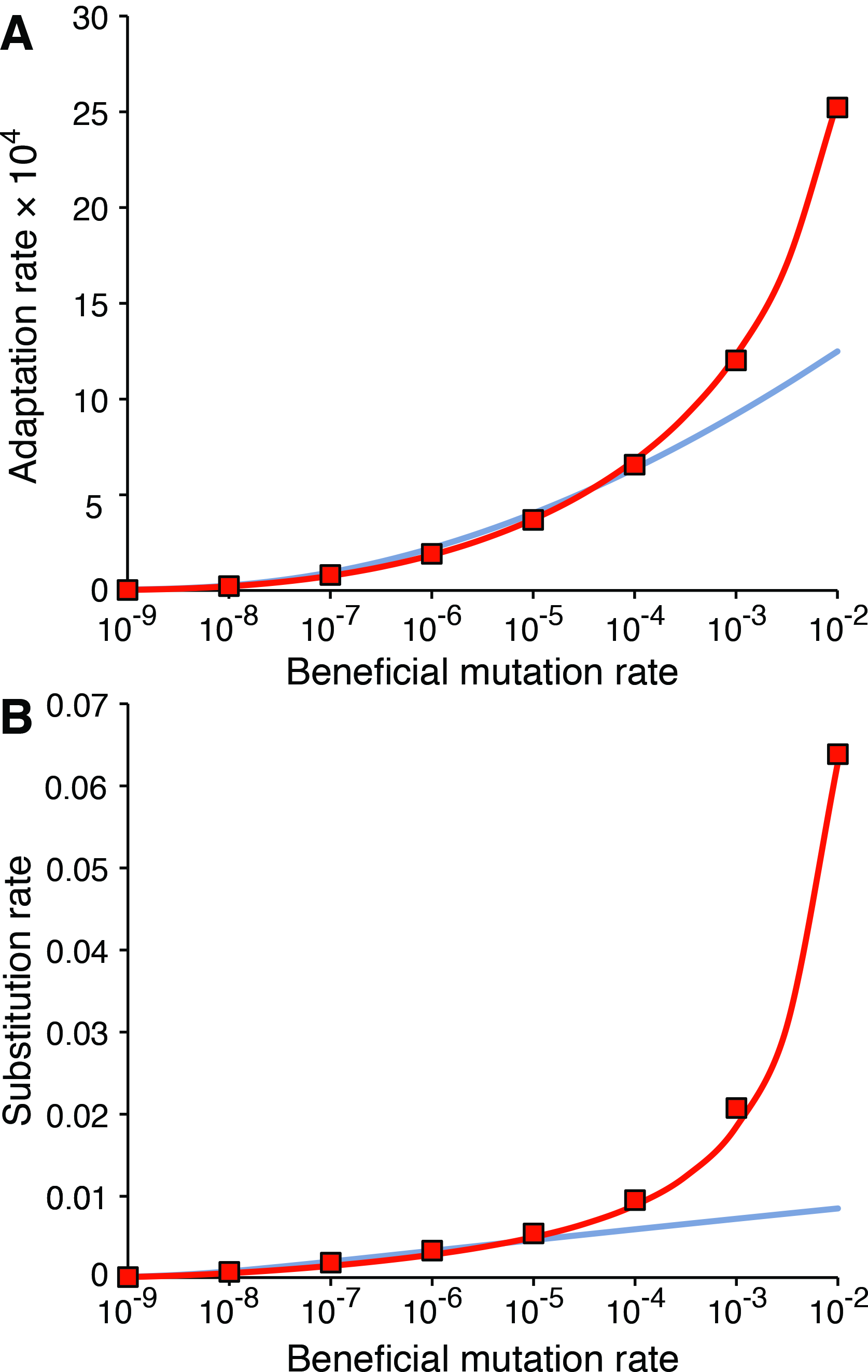
Lineage-interference theory accurately describes rates of adaptation and substitution in simulations over all mutation regimes. (A) Mean adaptation rate and (B) mean fixation rate for individual-based simulations (red symbols) compared to predictions from clonal-interference theory (blue curve) and lineage-interference theory (red curve). Population size *N* = 10^7^, mean of beneficial mutation effect distribution 1/*α* = 0.01.

Simulations were written in Python 2.7, using a Wright-Fisher model with discrete generations and infinite alleles. All distinct genotypes were tracked along with their corresponding frequencies, fitness values, and number of mutations. Reproduction was simulated by updating each genotype’s population size by drawing from a binomial distribution with *N*trials and 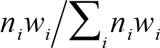 success probability in each trial, where *n_i_* and *w_i_* are the genotype’s population size and fitness, respectively. Mutation was simulated by randomly selecting a number of mutant individuals drawn from a binomial distribution with 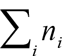 trials and 1−*e*^−*μ*^ success probability in each trial; 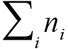 fluctuates slightly about *N*due to the stochastic implementation of reproduction, and 1−*e*^−*μ*^ is the Poisson probability that an individual receives at least one mutation. To assign a number of mutations to each mutant, numbers were iteratively drawn from the cumulative distribution function of a Poisson distribution with mean *μ*. Individuals receiving multiple mutations in a single generation were rare, even for the highest mutation rates considered here (up to 10^−2^). Each mutation was assigned with multiplicative effect on fitness, where the new fitness is *w_i_* (1 + *s*_mut_) and the effect size *s*_mut_ was drawn from an exponential distribution with mean 1/*α*.

We explored the evolutionary dynamics of lineages in the multiple-mutations regime using a graph-based implementation of our model (Fig. 1). Here, populations consisted of collections of unique genotypes, each of which was represented as a vertex. Rooted by the ancestral type, each mutation event created a new vertex, which was connected to its parent genotype by an edge. Along with the fitness effects of each mutation, vertices also stored the genotype’s abundance at each generation, allowing us to examine each mutation as it arose, spread, and either fixed or was lost. The abundance of a mutation at a given time was equal to the sum of all abundances in the subtree rooted at that vertex. After each generation, vertices were pruned if the corresponding genotype was extinct, had no extant descendants, and had not reached a threshold frequency. Reproduction and mutation were implemented as described above.

Adaptation rates from simulations were estimated by a least-squares linear fit to the population’s log-transformed mean fitness as a function of time in generations. One hundred runs of 100,000 generations each were performed for 10^−9^ ≤ *μ* ≤ 10^−3^, and 100 runs of 10,000 generations each were performed for *μ* = 10^−2^. To remove the transient effect of the initial accumulation of variation in fitness, the fit was started at the earliest point where all individuals had at least one mutation in at least half of the replicate runs; in the long simulations done here, this effect is very small. Substitution rates were measured by fitting a line to the minimum number of mutations (relative to the ancestor) in a population as a function of time in generations, starting at the same point as the fit to the adaptation rate.

## RESULTS

### Adaptation rate

The lineage-interference framework can be used to compute a population’s rate of fitness increase *v*. In order for a mutation to become fixed, it must occur, survive genetic drift, and survive interference. Following the logic of clonal-interference theory (GERRISH AND LENSKI 1998), we assume that these processes are independent: loss by drift occurs in the first few generations, whereas loss by interference typically occurs in later generations. Taking each of these processes in turn, we compute the expected initial advantage and expected fixation rate for new mutant lineages.

We assume that the effects of beneficial mutations *s*_mut_ are exponentially distributed, with probability density *α*exp(−*αs*_mut_), where 1/*α* is the mean effect. This distribution was chosen largely for mathematical convenience, because the exponential is a single-parameter distribution for which mutations of large effect are rare. Theoretical considerations (GILLESPIE 1984; ORR 2002; ORR 2003) and experimental evidence (IMHOF AND SCHLÖTTERER 2001; KASSEN AND BATAILLON 2006; MACLEAN AND BUCKLING 2009; FRENKEL *et al.* 2014) also support using an exponential distribution of beneficial fitness effects.

In the lineage-interference model, the mutant lineage’s advantage *s*_0_ depends on both the new beneficial effect and the background fitness. As was done previously (ROUZINE AND COFFIN 2005; DESAI AND FISHER 2007; NEHER *et al.* 2010; GOOD *et al.* 2012), we approximate the distribution of background fitness values as a Gaussian with variance *σ*^2^ According to Fisher’s Fundamental Theorem of Natural Selection (FISHER 1930), a population’s rate of fitness increase *v* is equal to the genetic variance in fitness *σ*^2^ This relation creates a self-consistency condition that constrains the value of the adaptation rate, which we will use to calculate solutions for *v*.

The mutant lineage’s initial advantage over the population mean fitness is therefore given by the sum of two random variables: a Gaussian with *σ*^2^ = *v* representing background fitness values, and an exponential with mean 1/*α* representing the fitness effects of new mutations. The distribution of their sum is given by their convolution, which is an exponentially modified Gaussian:

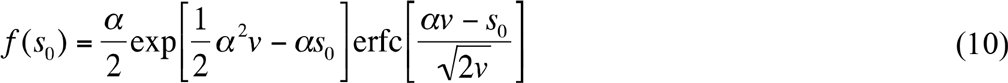

where erfc is the complementary error function. For small *v*, there is negligible effect from the background, and this distribution reduces to an exponential.

After a new beneficial mutation occurs, it must survive genetic drift while it remains at low frequencies. In a large constant-size population where each individual has a Poisson distributed number of offspring, the probability that a mutation with small effect size *s*_0_ survives drift is given by Haldane’s formula (HALDANE 1927):

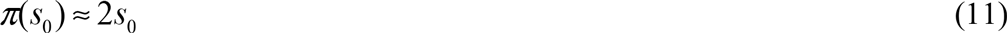

Finally, in order for a mutation to become fixed, it must survive interference, which we have shown requires *s*_0_ > *s*_crit_, where *s*_crit_ is given by Eq. (9). For multiplicative fitness, the expected contribution to adaptation of a fixed beneficial mutant is:

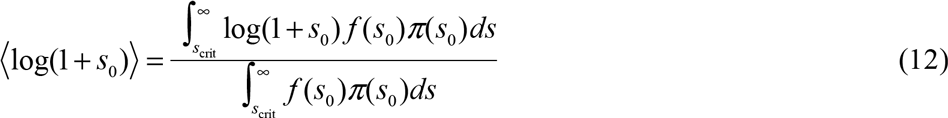

and the rate at which mutant lineages arise that eventually fix is:

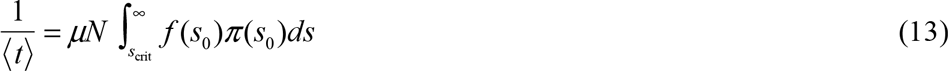

where *μ* is the beneficial mutation rate per individual per generation. The adaptation rate is then:

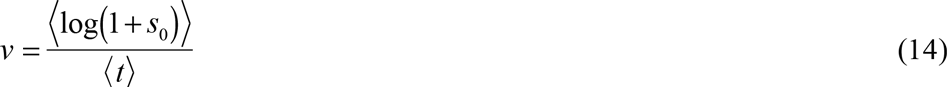

where *v* is the rate of exponential increase in fitness. The right side of Eq. (14) depends on input parameters *μ*, *N*, and *α* as well as on *v* itself. Therefore, Eq. (14) cannot be solved explicitly for *v*, and solutions must be computed numerically. These solutions are compared to individual-based simulations (see Methods). Fig. 4A shows that the lineage-interference model accurately describes adaptation rates in all selection regimes: the low *Nμ* periodic-selection regime (*Nμ* ≪ 1/log(*α*/*N*, i.e., *μ* < 10^−8^ in the simulations) (DESAI AND FISHER 2007), the moderate *Nμ* clonal-interference regime (log(*Nμ*) ≪ 2log(1/*μα*), i.e. 10^−8^ < *μ* < 10^−4^ in the simulations) (GOOD *et al*. 2012), and the high *Nμ* multiple-mutations regime where clonal-interference theory fails (*μ* > 10^−4^ in the simulations).

We can simplify this expression for *v* in certain parameter regimes. if the expected fixed advantage is not too large (〈*s*_0_〉 ≪ 1), and if the population size is not too small 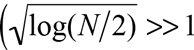 and 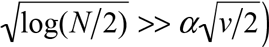, then *v* can be expressed as a simpler transcendental equation:

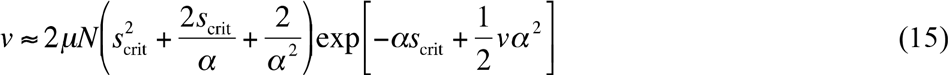

This approximation is equivalent to a solution obtained by Good *etal*. (GOOD *etal*. 2012), who used branching-process theory to compute the adaptation rate. A comparison of that work to the lineage-interference model can be found in the Discussion section below.

### Substitution rate

By considering a new mutant’s fitness that accounts for its genetic background as well as its individual effect, the lineage-interference model provides a convenient framework for assessing a mutant lineage’s fate. However, this treatment complicates the identification of individual fixed mutations because any particular mutant lineage might include additional mutations on the background that have yet to fix, which cannot be directly quantified. Therefore, we must probabilistically estimate the expected number of currently unfixed mutations on the background of a mutation that will likely become fixed.

Consider a new mutant lineage with total advantage *s*_0_, which is composed of a mutational effect *xs*_0_ and background effect (1−*x*)*s*_0_. Because the two distributions are independent, the non-normalized joint probability density is equal to their product:

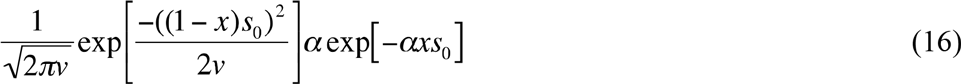

Normalizing this probability density over the interval 0 < *x* < ∞ (to infinity because the background advantage can be negative), we have:

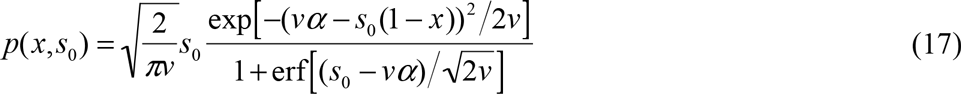

where erf is the error function. Eq. (17) is the probability density of the fraction *x* of the total mutant advantage *s*_0_ that is attributed to the effect of the mutation itself. The expected fraction ⟨*x*⟩ attributed to a fixed mutational effect is:

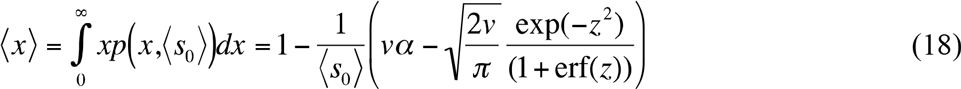

where 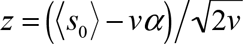. Because any beneficial mutation may in principle be called the “focal” mutation that defines a lineage, the expected number of mutations contained in a fixed lineage is simply 1/⟨*x*⟩. Then, the substitution rate for new mutations is:

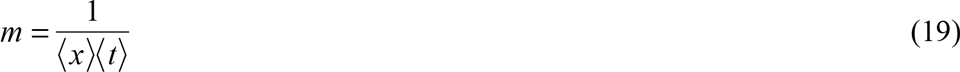

As Fig. 4B shows, this predicted substitution rate agrees closely with results from individual-based simulations (see Methods).

## DISCUSSION

We have described a comprehensive conceptual model of competing lineages in asexual populations. The framework for this model is based on four simple, key observations.

1. The likely fate of any given beneficial mutation is determined by a combination of its effect size and its background fitness.
2. Background fitness effects are directly tied to the overall adaptation rate by Fisher’s Fundamental Theorem.
3. The adaptation rate of any particular lineage is proportional to its frequency in the population at any given time, which implies a minimum required initial advantage for eventual fixation of mutants.
4. Any mutation may be considered one that defines a lineage, which allows for estimating the number of beneficial mutations that a typical successful lineage accumulates on its way to fixation.

This “lineage-interference” model accurately describes the rates of both adaptation and substitution over all selection regimes, including the multiple-mutations regime where clonal-interference theory breaks down (GERRISH AND LENSKI 1998). We have confirmed using simulations that the model works for mutation supply rates up to *Nμ* = 10^5^, which is the highest that was computationally feasible; this confirmation even includes high values of *μ* on the order of 1/*α*, where the Gaussian approximation of background fitness values breaks down. Lineage-interference theory also provides a mathematically simple interpretation of interference, which is realized by simply adding up all mutants above a particular threshold (see Eqs. (12) and (13)).

Until recent years, theoretical progress on the multiple-mutations regime had been limited to constant effect-size models, using the “traveling wave” framework (DESAI AND FISHER 2007; ROUZINE *et al.* 2008; NEHER *et al.* 2010). This work provided valuable insights into the transition from stochastic to deterministic fates of new mutants, and the qualitative behavior of adaptation rates with increasing *Nμ*. However, in general real mutation frequency dynamics involves interference among mutations of different effect sizes, which requires incorporating a continuous distribution of effects into the theory.

To our knowledge, there is currently only one other model of adaptation in the multiple-mutations regime that uses a continuous distribution of mutational effects. Good *et al.* (GOOD *et al.* 2012) approximate a sharp transition in the fixation probability for mutations above a particular effect size *x_c_*, which is given by a transcendental integro-differential equation from branching process theory. The sharp transition they describe is directly analogous to the critical threshold *s*_crit_ in our model—in fact, they obtain an identical large-*N* approximation of the adaptation rate, where their *x_c_* is equivalent to our *s*_crit_. However, by framing the fate of each beneficial mutation directly in the context of its background, we have been able to compute a simple, closed-form solution for the critical advantage 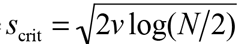 (see Eqs. (6-9)), which is independent of any assumptions about rates, effect sizes, and distributions of mutations.

Conceptualizing an adapting population as a collection of competing lineages can also help us understand the transition to the multiple-mutations regime at high *Nμ*. As the mutation supply rate increases, multiple mutations occur on the same background before any single one can fix, typically resulting in a better competitor than any single mutant (Fig. 1). This breaks the adaptation “speed limit” of clonal-interference theory, which only considers the competition between individual mutations. Lineage interference accounts for multiple mutations, and it accurately predicts outcomes on both sides of the regime transition.

The transition to the multiple-mutations regime can be illustrated by the typical number of mutations that comprise a successful lineage (Fig. 5A). At the point where the clonal-interference model breaks down (at *μ* > 10^−4^ in the parameterization shown in Figs. 4 and 5), a successful lineage already contains ~2 beneficial mutations. This point can be considered the beginning of “nesting” selective sweeps, where a second strong mutation occurs just before the previous one fixes, barely helping to make the first one successful. At even higher *Nμ*, lineages must accumulate additional beneficial mutations earlier and earlier in their respective histories in order to remain competitive. Eventually, sweeps of fixing mutations become more and more highly “nested” until they resemble “cohorts” of mutations sweeping roughly in lockstep. The lineage-interference model may thus help to quantitatively understand the cohorts of multiple beneficial mutations that have been observed in some evolution experiments (LANG *et al*. 2013; MADDAMSETTI *et al*. 2015).

**Figure 5.**
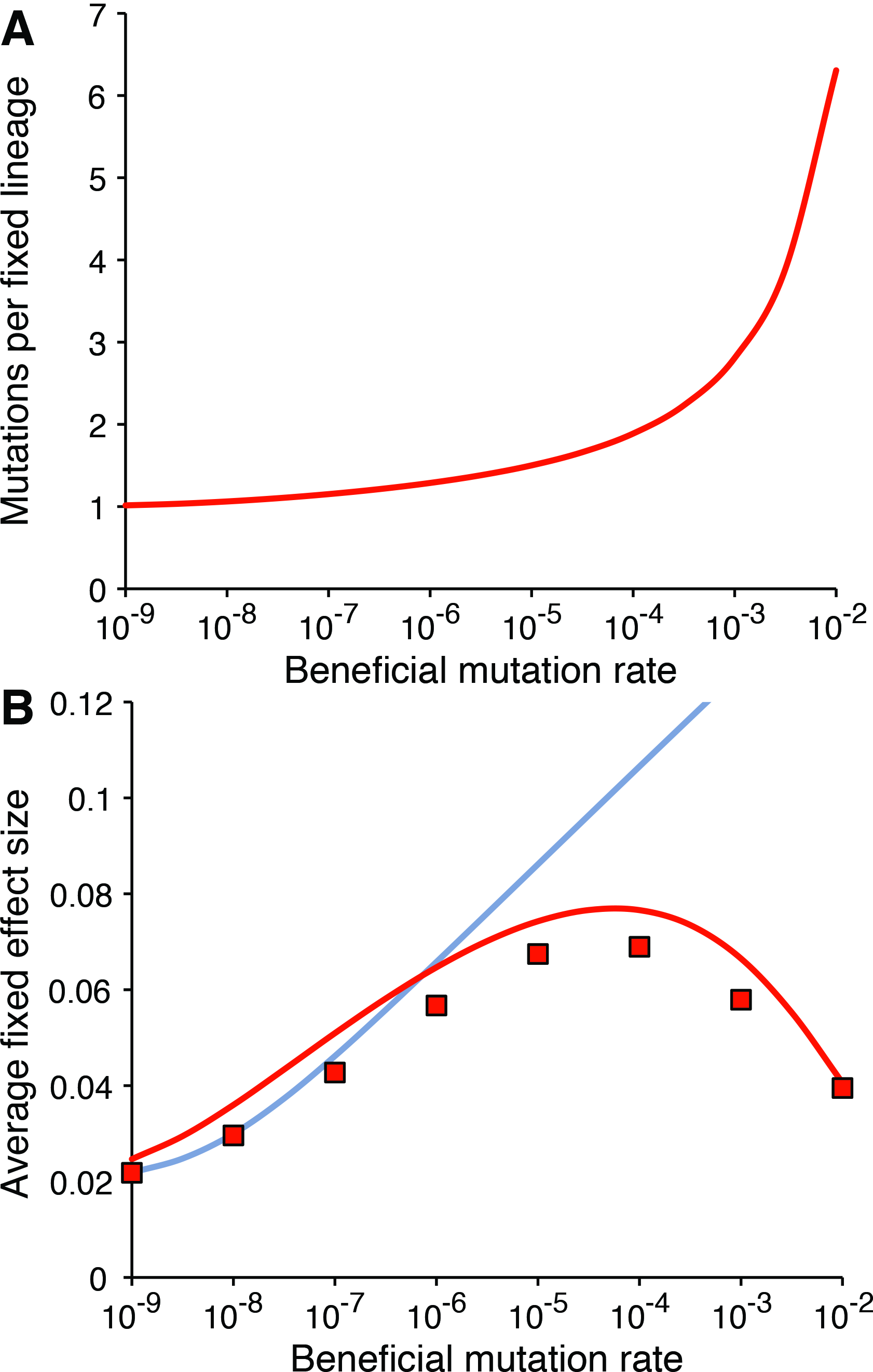
Fixed lineages comprise multiple smaller-effect mutations in the multiple-mutations regime. (A) Number of mutations contained in an average fixed lineage predicted by the lineage-interference model, as a function of the beneficial mutation rate. (B) Mean effect size of a fixed beneficial mutation in simulations (red symbols), and the same values predicted by the lineage-interference (red curve) and clonal-interference (blue curve) models as a function of the beneficial mutation rate. Population size *N* = 10^7^, mean of beneficial mutation effect distribution 1/*α* = 0.01.

The effects of the multiple-mutations regime can also be seen in the average effect size of a fixed beneficial mutation (Fig. 5B). In the clonal-interference regime, higher beneficial mutation rates result in fixed mutations of larger effect size, as the competition between individual mutations becomes fiercer. But at some point, two individual mutations with smaller beneficial effects are more likely to co-occur in a lineage than any single large effect mutation. As a result, increasing the beneficial mutation rate above some level leads counter-intuitively to the fixation of smaller effect mutations.

The largest anomaly generated by approximation in the lineage-interference model is also well illustrated in Fig. 5B: the model slightly overestimates (by ~15%) the average effect size of fixed mutations. Furthermore, predictions of the substitution rate are underestimated by roughly the same amount (see Fig. 4B). These deviations result from the assumption that *s*_crit_ represents a hard threshold required for fixation of a mutant lineage. More accurately, *s*_crit_ should be treated as a characteristic value that describes the likelihood of fixation, where some lineages below the threshold become fixed and some above the threshold do not. As a result, the model described here predicts the fixation of slightly fewer mutations of slightly larger effect than those that actually become fixed. Although this approximation does not affect estimates of the overall adaptation rate, it may be possible in future work to relax this assumption to obtain even more accurate descriptions of substitution dynamics.

The lineage-interference model also reveals features of the intermediate *Nμ* regime that the clonal-interference theory does not capture. To isolate these effects, Fig. 6 shows adaptation rates from the lineage-interference model and from simulations, both plotted relative to the clonal-interference model. shown this way, simulated adaptation is clearly slower (by ~15% at *Nμ* = 1) than predicted by clonal-interference theory, precisely in the regime that it is meant to describe. In comparison, the lineage-interference model accurately captures this deviation. This deviation indicates that there are systematic effects of multiple mutations even outside the multiple-mutations regime: specifically, some beneficial mutations can be “wasted” by virtue of their falling on backgrounds of below-average fitness. When interference is not strong, the same mutation would gain negligible advantage by occurring on an above-average fitness background, because it is likely to become fixed anyway by virtue of being “good enough.” The net result is that multiple-mutation effects actually impede adaptation in the intermediate *Nμ* regime. This effect could be important when interpreting the consequences of mutator alleles in evolving populations (TADDEI *et al.* 1997; SHAVER *et al.* 2002; RAYNES *et al.* 2012; WIELGOSS *et al.* 2012): the potential evolutionary advantage gained by increasing the beneficial mutation rate in this regime could be significantly lessened.

**Figure 6.**
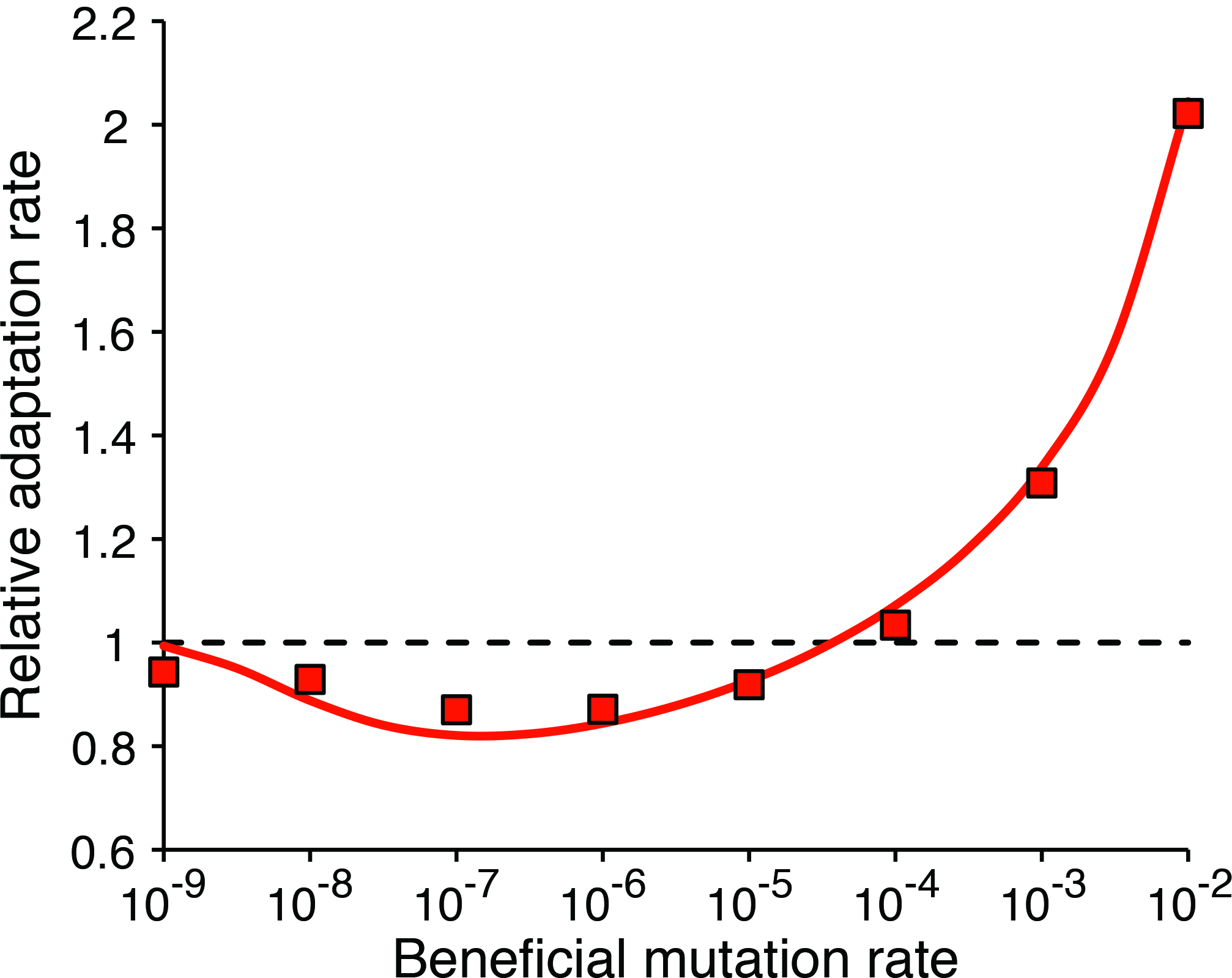
Multiple-mutation effects impede adaptation in the clonal-interference regime. The red curve shows the mean adaptation rate predicted by lineage-interference theory divided by the corresponding prediction from clonal-interference theory. The red symbols are simulation results divided by the clonal-interference prediction. Population size *N* = 10^7^, mean of beneficial mutation effect distribution 1/*α* = 0.01.

In practice, it is difficult to experimentally test models of adaptation that involve many fixed mutations, because it may require thousands of generations to observe enough selective sweeps to accurately measure adaptation rates. For large population sizes where multiple mutations are most relevant, sweep times are even longer. In Lenski’s long-term evolution experiment (LTEE) with *Escherichia coli*, the adaptation rate was analyzed using a clonal-interference framework (Wiser *et al.* 2013). In that study, the upper 95% confidence limit of *Nμ* was placed at ~2 × 10^3^ (for populations that had not evolved hypermutability), which would put it barely in the multiple-mutations regime (*Nμ* > 10^3^ according to Fig. 4A). Each population in this experiment is given 10 mL of a minimal medium with a low resource concentration and experiences daily 100-fold dilutions; larger laboratory populations (e.g., in rich media, in larger volumes, or in continuous culture without bottlenecks) would have larger effective population sizes and therefore might be well within the multiple-mutations regime.

In order to compare fitness data from laboratory experiments to the lineage-interference model, future research will need to incorporate some quantitative description of epistasis into the theory. Making that challenge more tractable, previous work in several systems has shown that typical populations exhibit a general trend of diminishing-returns epistasis (MACLEAN *et al.* 2010; CHOU *et al.* 2011; KHAN *et al.* 2011; KRYAZHIMSKIY *et al.* 2014), which can be parameterized in a simple way that accurately describes the dynamics of mean fitness (WISER *et al.* 2013). Diminishing-returns epistasis means that the combined effect of beneficial mutations is less than their sum, and lower fitness backgrounds have access to beneficial mutations with larger effect sizes. This form of epistasis would alter the lineage-interference model in three distinct ways. First, the adaptation rate of each lineage would depend not only on its frequency, but also on its mean fitness. Second, *f* (*s*_0_) would have to be adjusted to account for the non-additivity of fitness effects. Third, *v* would need to be calculated sequentially for each lineage fixation, because the mean of the distribution of available beneficial effects would decline after each fixation; this third effect was previously incorporated into the clonal-interference framework (WISER *et al.* 2013). It remains to be seen whether these issues can be formally incorporated into the lineage-interference framework, or whether individual-based simulations will have to suffice when dealing with the complications introduced by epistasis.

In conclusion, the lineage-interference model provides a valuable conceptual framework for understanding the likely fates of beneficial mutations and the increased adaptation rate in the transition to the multiple-mutations regime. We have demonstrated that this model accurately predicts a population’s adaptation rate based on a simple integral over all mutant lineages with a fitness advantage above a critical threshold that is proportional to 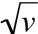. This prediction indicates that multiple-mutation effects occur even in the clonal-interference regime, and that they impede adaptation. We have described a probabilistic framework for identifying the likely number of beneficial mutations present in each fixing lineage, which illustrates quantitatively how mutations “team up” when the mutation supply rate is high. We expect this framework will help elucidate the cause of cohorts of beneficial mutations sweeping together, and will be a useful tool for assessing how rapidly large populations adapt to their environments.

## ACKNOWLEDGMENTS

This work was supported by the National Science Foundation BEACON Center for the Study of Evolution in Action (cooperative agreement DBI-0939454) and by the National Science Foundation (DEB-1451740 to R.E.L.). We are grateful to the Helen Riaboff Whiteley Center and Friday Harbor Laboratories for providing accommodations and working space for our collaboration.

